# The contribution of ancient admixture to reproductive isolation between European sea bass lineages

**DOI:** 10.1101/641829

**Authors:** Maud Duranton, François Allal, Sophie Valière, Olivier Bouchez, François Bonhomme, Pierre-Alexandre Gagnaire

## Abstract

Understanding how new species arise through the progressive establishment of reproductive isolation barriers between diverging populations is a major goal in Evolutionary Biology. One important result of speciation genomics studies is that the genomic regions involved in reproductive isolation frequently harbor anciently diverged haplotypes that predate the reconstructed history of species divergence. The possible origins of these old alleles remain highly debated, since they relate to contrasted mechanisms of speciation that are not fully understood yet. In the European sea bass (*Dicentrarchus labrax*), the genomic regions involved in reproductive isolation between Atlantic and Mediterranean lineages are enriched for anciently diverged alleles of unknown origin. Here, we used haplotype-resolved whole-genome sequences to test whether divergent haplotypes could have originated from a closely related species, the spotted sea bass (*Dicentrarchus punctatus*). We found that an ancient admixture event between *D. labrax* and *D. punctatus* is responsible for the presence of shared derived alleles that segregate at low frequencies in both lineages of *D. labrax*. An exception to this was found within regions involved in reproductive isolation between the two *D. labrax* lineages. In those regions, archaic tracts originating from *D. punctatus* locally reached high frequencies or even fixation in Atlantic genomes but were almost absent in the Mediterranean. We showed that the ancient admixture event most likely occurred between *D. punctatus* and the *D. labrax* Atlantic lineage, while Atlantic and Mediterranean *D. labrax* lineages were experiencing allopatric isolation. Our results suggest that local adaptive introgression and/or the resolution of genomic conflicts provoked by ancient admixture have probably participated to the establishment of reproductive isolation between the two *D. labrax* lineages.

**Author summary:** Speciation is often viewed as a progressive accumulation of reproductive isolation barriers between two diverging lineages through the time. When initiated, the speciation process may however take different routes, sometimes leading to the erosion of an established species barrier or to the acquisition of new speciation genes transferred from another species boundary. Here, we describe such a case in the European sea bass. This marine fish species has split 300,000 years ago into an Atlantic and a Mediterranean lineage, which remained partially reproductively isolated after experiencing postglacial secondary contact. For unknown reasons, genomic regions involved in reproductive isolation between lineages have started to diverge well before the split. We here show that diverged alleles were acquired by the Atlantic lineage from an ancient event of admixture with a parapatric sister species about 80,000 years ago. Introgressed foreign alleles that were locally driven to high frequencies in the Atlantic have subsequently resisted to introgression within the Mediterranean during the postglacial secondary contact, thus contributing to increased reproductive isolation between two sea bass lineages. These results support the view that reproductive isolation barriers can evolve via reticulate gene flow across multiple species boundaries.

## Introduction

Speciation is the evolutionary process that leads to the emergence of new species through the progressive establishment of Reproductive Isolation (RI) barriers between diverging populations (1). Identifying those barriers and understanding the eco-evolutionary context in which they evolved has been at the core of the speciation genetics research program (2,3). Over the last decade, progresses in sequencing technologies have allowed to gain important insights into the genetic basis of reproductive isolation barriers through the study of genome-wide differentiation/divergence patterns between closely related species (4–8). An important result of speciation genomics studies was that the age of the alleles located within genomic regions involved in RI is often much older than the average coalescent time computed across the whole genome. This finding indicates that the regions involved in RI tend to be enriched for anciently diverged haplotypes. An example of this comes from the fixed chromosomal inversions involved in RI between *Drosophila pseudoobscura* and *D. persimilis*, which show higher divergence than collinear regions of the genome (9). Another case is provided by the large genomic regions of ancient ancestry that have been found across the threespine stickleback’s genome, which are involved in RI between marine and freshwater populations (10,11). A third example, among others (see <u>Marques *et al.* (2019)</u> for a review), was described in Darwin’s finches, whereby genomic regions showing increased divergence in several species pairs also display anciently diverged haplogroups that originated before the species splits (13).

Different hypotheses can explain the origin and the maintenance of these highly divergent haplotypes. First, polymorphism has possibly been maintained over the long term in the ancestral population before being differentially sorted between the descendant lineages (14). This hypothesis has been proposed to explain the excess of haplotype divergence in the aforementioned examples (9,10,13). One mechanism that may explain the long-term maintenance of polymorphism is ancestral population structure, that is, subdivision owing to barriers to gene flow in the ancestral population (15). In addition to demography, balancing selection due to either frequency-dependent selection, heterozygote advantage (overdominance) or heterogeneous selection in space or time (16) can also promote the maintenance of ancient polymorphisms. For instance, in Darwin’s finches, balancing selection has been proposed to explain the maintenance of divergent haplogroups associated with beak shape, due to the selective advantage of rare beak morphologies, or changing environmental conditions inducing heterogeneous selection (13). An alternative explanation to the presence of anciently diverged alleles is admixture with a divergent lineage. Contemporary hybridization has long been recognized as a common phenomenon in plants and animals (17,18), and cases of ancient admixture are increasingly detected by genomic studies. One emblematic example is past admixture between modern humans and two extinct archaic hominin lineages, Neanderthal and Denisova (19– 21). More recently, ancient introgression from the extinct cave bear has also been detected in the genomes of living brown bears (22). Therefore, past admixture is increasingly recognized as a source of anciently diverged alleles in contemporary genomes.

Understanding why and how divergent haplogroups tend to disproportionately contribute to the buildup of RI between nascent species remains, however, highly challenging. First, because retention of ancestral polymorphism and past admixture are notoriously difficult to distinguish and not mutually exclusive hypotheses to explain the presence of anciently diverged alleles (23–27). Furthermore, identifying the genomic regions that resist introgression is still a major obstacle to the detection of RI loci (28). These tasks are now facilitated by the direct assessment of local ancestry along individual genome sequences (29,30), thus paving the way for assessing the role of ancient admixture in speciation. Here, we use new haplotype-resolved whole-genome sequences to delineate the regions involved in RI between European sea bass lineages and understand the origin of the divergent haplogroups they contain.

The European sea bass (*Dicentrarchus labrax*) is a marine fish subdivided into two glacial lineages, which currently correspond to Atlantic and Mediterranean populations (31). These two lineages have diverged in allopatry for *c.a.* 300,000 years before experiencing a secondary contact since the last glacial retreat (32). Postglacial gene flow between the two lineages is strongly asymmetrical, mostly occurring from the Atlantic to the Mediterranean genetic background (32). This resulted in a spatial introgression gradient within the Mediterranean Sea, illustrated by a more than twofold higher Atlantic ancestry in the western (31%) compared to the eastern (13%) Mediterranean population (30). A detailed analysis of local ancestry tracts across Mediterranean and Atlantic sea bass genomes has provided direct evidence for highly heterogeneous rates of gene flow along most chromosomes (Duranton *et al.* 2018). This mosaic introgression pattern was attributed to the effect of multiple small effect RI loci mainly located in low-recombining regions that present particularly high values of nucleotidic divergence (*d*_XY_). It is generally assumed that increased *d*_XY_ indicates the presence of haplotypes that started to diverge earlier than the rest of the genome. However, regions of increased divergence may simply have resisted gene flow during secondary contact, while haplotypes in the remainder of the genome got rejuvenated due to recombination. This later hypothesis, however has been rejected in the European sea bass using simulations accounting for both background selection and selection against introgressed tracts (30). Therefore, anciently diverged alleles are unlikely to have evolved within the 300,000 years divergence history inferred from genome-wide polymorphism data and are thus older. In the present study, we use new haplotype-resolved whole-genome sequences to accurately delineate regions involved in RI and investigate the mechanisms underlying their excess of divergence. We specifically test for past admixture with a closely related species using a new genome sequence from the parapatrically distributed spotted sea bass (*Dicentrarchus punctatus*). Our results show that gene flow occurred between *D. punctatus* and the Atlantic lineage of *D. labrax* about 80,000 years ago, resulting in a low background ancestry from *D. punctatus* in contemporary *D. labrax* genomes. By contrast, genomic regions involved in RI between the two *D. labrax* lineages generally display high frequencies of haplotypes derived from *D. punctatus* in the Atlantic, while these archaic tracts remain rare in the Mediterranean. This suggests that ancient admixture has played an important role in the evolution of RI between Atlantic and Mediterranean sea bass lineages, consistently with predictions from models of local adaptive introgression and selection against genetic incompatibilities.

## Results

### Phylogenomic analysis

We reconstructed the genetic relationships among the three Moronid species used in our study: the striped bass (*Morone saxatilis*), the spotted sea bass (*Dicentrarchus punctatus*) and the European sea bass (*Dicentrarchus labrax*), which is further subdivided into two partially reproductively isolated populations: the Atlantic and Mediterranean sea bass lineages. All of the 3,329 maximum-likelihood phylogenetic trees generated in non-overlapping 50kb windows showed the same topology, corresponding to the expected species tree (Figure 1A). However, when similar reconstructions were performed in 2 kb windows, 4.6% of conflicting genealogies were found with an excess of trees in which *D. punctatus* grouped with the Atlantic (2.87%) versus with the Mediterranean (1.68%) *D. labrax* lineage (Supplementary Figure 3). The relative branch lengths of the species tree largely reflected the mean nucleotide divergence (*d*_XY_) measured between each pair of four species/lineages (Figure 1B). We found 4.5% of absolute sequence divergence between the outgroup *M. saxatilis* and the two *Dicentrarchus* species. Divergence between *D. labrax* and *D. punctatus* (0.55%) was more than five times higher than divergence between Atlantic and Mediterranean *D. labrax* lineages (0.1%), consistently with previous estimates (30,32). We found a slightly higher divergence between *D. punctatus* and the eastern Mediterranean (0.56%) compared to the Atlantic *D. labrax* lineage (0.53%). Within *D. labrax*, divergence to the Atlantic population was higher for the eastern (0.1%) compared to the western Mediterranean population (0.09%) (consistent with the PCA, Supplementary Figure 2), as expected due to gene flow between Atlantic and Mediterranean *D. labrax* lineages (30,32).

**Figure 1.**
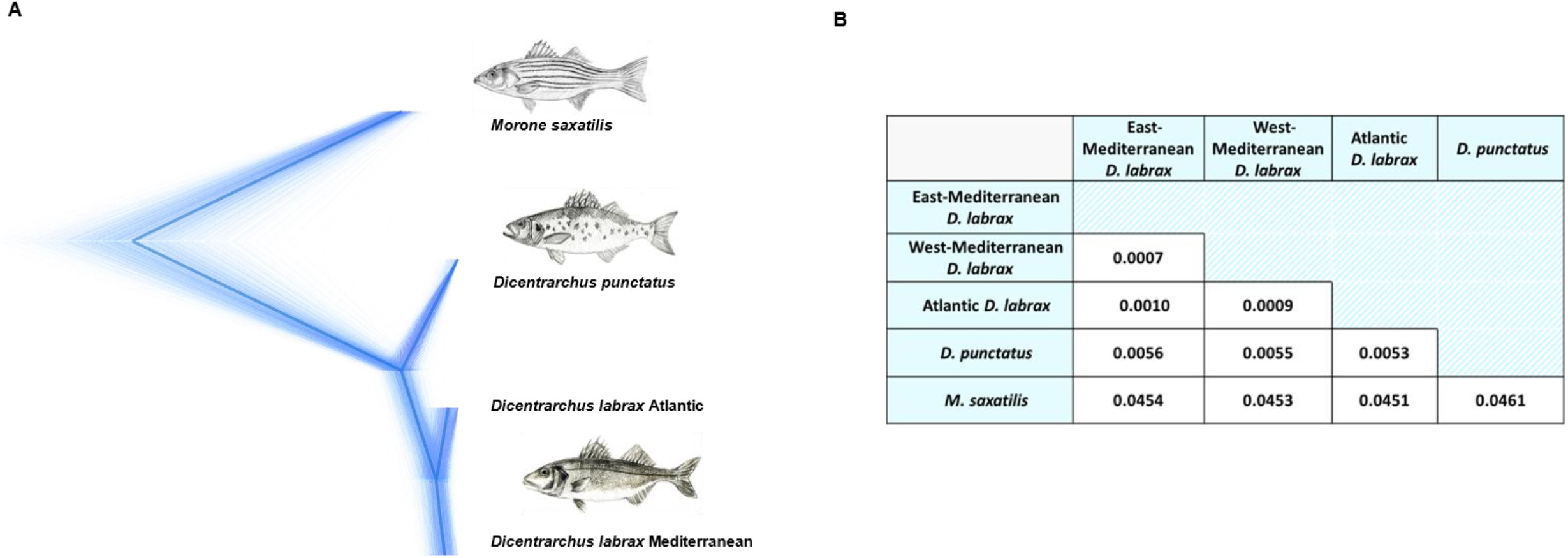
Phylogenetic relationships between *D. labrax* lineages, *D. punctatus* and the outgroup species *M. saxatilis*. **A.** Concordance of 3,329 maximum-likelihood trees reconstructed in non-overlapping 50 kb windows along the genome (thick lines) and superimposed to the consensus species tree (bold line) using DensiTree v2.2.5 (bouckaert and heled 2014). Only one haplome from each species/lineage was used for tree reconstruction. Discordant trees that disproportionately grouped the Atlantic *D. labrax* lineage with *D. punctatus* were observed at a more local scale using 2 kb windows (Supplementary Figure 3). **B.** Genome-wide average pairwise sequence divergence between species/lineages measured by *d*_xy_ using the same individual haplotypes as for the phylogenetic relationships.

### Test for foreign introgression within *D. labrax*

Chromosomal patterns of absolute sequence divergence (*d*_XY_) between the Atlantic and Mediterranean lineages of *D. labrax* (Fig 2A and Supplementary Figure 4A) showed highly heterogeneous divergence along the genome, as reported in previous studies (30,32). To determine if local excesses of *d*_XY_ can be explained by past admixture with another lineage, we first looked for gene flow between *D. labrax* lineages and *D. punctatus* using the ABBA-BABA test. Some genomic regions showed particularly high values of the *f*_D_ statistics, thus reflecting locally elevated ancestry from *D. punctatus* within the *D. labrax* Atlantic lineage (Figure 2B and Supplementary Figure 4B red curve). By contrast, when the *f*_D_ statistics was used to measure local *D. punctatus* ancestry within *D. labrax* Mediterranean populations, low and relatively homogeneous introgression patterns were found across the entire genome (Figure 2B and Supplementary Figure 4B blue and green curves). This finding thus indicates highly heterogeneous introgression of spotted sea bass alleles within the Atlantic *D. labrax* lineage, and comparatively lower introgression within the Mediterranean lineage.

**Figure 2.**
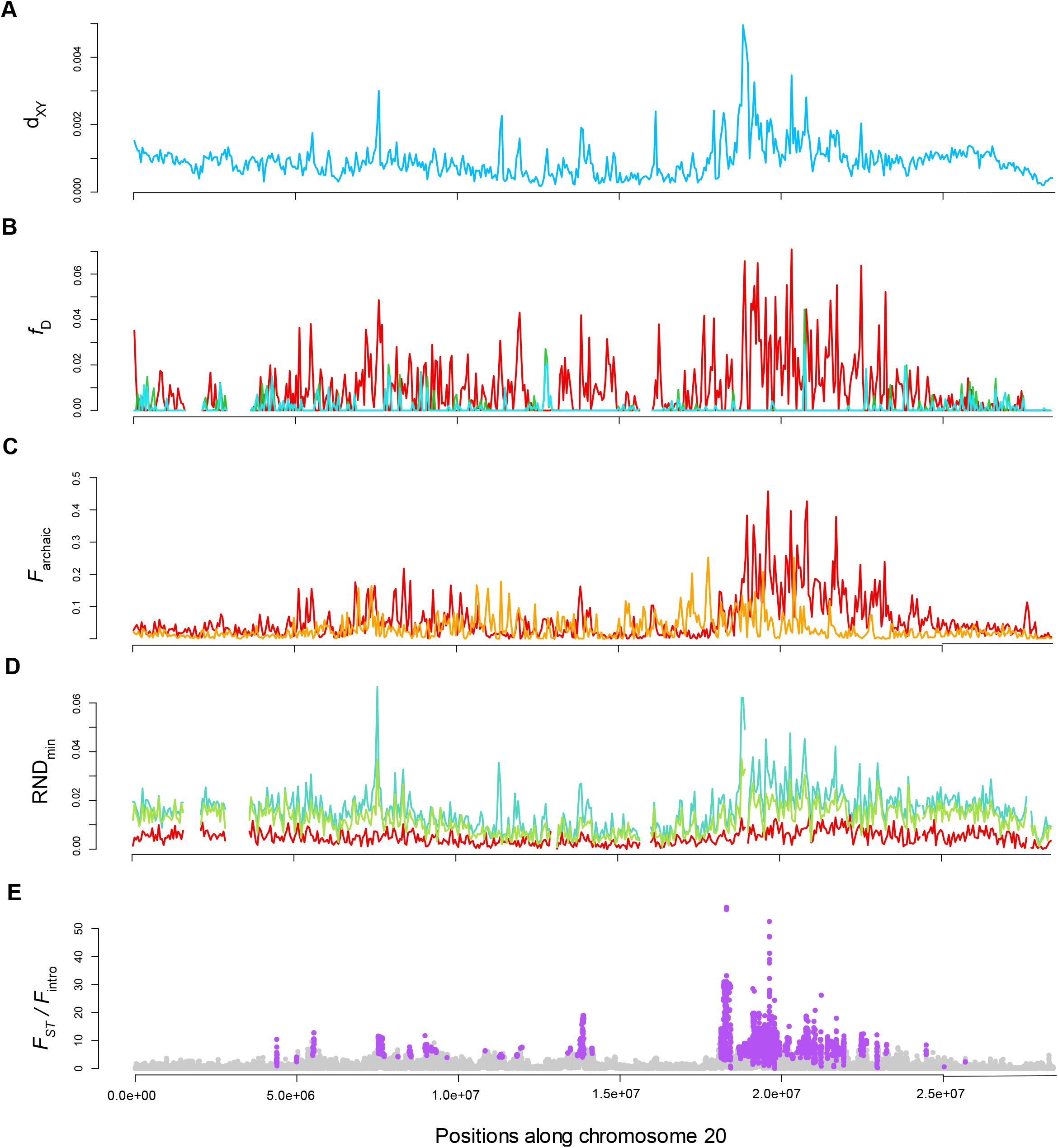
Divergence and introgression statistics measured in non-overlapping 50 kb windows along chromosome 20. **A.** *d*_XY_ measured between the Atlantic and Mediterranean (including eastern and western population) *D. labrax* lineages. **B.** *f*_D_ statistics measured using (((MED, AT), PUN), SAX) in red, (((AT, WEM), PUN), SAX) in green, and (((ATL, SEM), PUN), SAX) in blue. **C.** Fraction of archaic introgressed tracts (*F*_archaic_) inferred in the eastern Mediterranean and Atlantic populations of *D. labrax*. **D.** RND_min_ measured between *D. punctatus* and *D. labrax* Atlantic (red), western (green) and eastern (blue) Mediterranean populations. **E.** *F*_ST_ between the Atlantic and western Mediterranean population of *D. labrax* divided by the fraction of Atlantic tracts introgressed into the western Mediterranean genomes for each SNP along the chromosome. Purple points show SNPs with significant associations to reproductive isolation after applying FDR correction to the probabilities determined with the HMM approach.

We also searched for the presence of archaic introgressed tracts in *D. labrax* genomes. A relatively low fraction of archaic tracts (*F*_archaic_) was found along the genome in both Atlantic (4.85% in non-RI islands) and Mediterranean (2.73% in non-RI islands) *D. labrax* individuals (Figure 2C and Supplementary Figure 4C). In some regions, however, *F*_archaic_ was particularly high in the Atlantic (i.e. >30%) compared to the Mediterranean lineage. Interestingly, those regions also presented the highest *f*_D_ values (Figure 2B red curve), and there was a highly significant positive correlation between *f*_D_ and *F*_archaic_ in Atlantic *D. labrax* genomes (Spearman’s rho = 0.281***). These results thus support the hypothesis that the detected archaic segments that locally reach high frequencies in some regions of Atlantic *D. labrax* genomes have been inherited from *D. punctatus* at some time in the past. Furthermore, regions of particularly increased *D. punctatus* ancestry also showed the highest absolute divergence values between Atlantic and Mediterranean *D. labrax* lineages, with positive genome-wide correlations being found with *d*_XY_ for both *f*_D_ (Spearman’s rho = 0.281***) and *F*_archaic_ (Spearman’s rho = 0.531***). Lastly, we used the RND_min_ statistics to detect chromosomal variations in ancient introgression. Values of RND_min_ measured between *D. punctatus* and the Atlantic *D. labrax* lineage were low and relatively constant along chromosomes (Figure 2D and 4D red curves), indicating widespread (although locally rare) introgression across the genome. By contrast, RND_min_ was higher and highly variable when measured with the Mediterranean *D. labrax* populations (Figure 2D and 4D blue and green curves), indicating that introgression from *D. punctatus* is absent or nearly absent in some genomic regions of the Mediterranean lineage. These regions, that seem resistant to *D. punctatus* introgression in Mediterranean *D. labrax* genomes, also showed elevated values of *F*_archaic_ (genome-wide Spearman’s rho = 0.472***) and *f*_D_ (genome wide spearman’s rho = 0.223***) in Atlantic genomes, along with increased *d*_XY_ between Atlantic and Mediterranean *D. labrax* lineages (genome-wide Spearman’s rho = 0.717***). These results thus indicate the existence of outlying patterns of *D. punctatus* ancestry in the most divergent genomic regions between *D. labrax* lineages, due to respectively increased and decreased frequencies of anciently introgressed tracts in the Atlantic and Mediterranean lineages, compared to the background level.

Finally, our HMM approach allowed categorizing 70,738 SNP that are likely associated with RI islands between the two *D. labrax* lineages (Figure 2E and Supplementary Figure 4E). We found a good concordance between the positions of RI islands identified with the SNP and window-based methods, although the former allowed us to detect narrower RI-associated regions with a higher resolution (Supplementary Figure 5C and F). As expected, all these regions displayed increased levels of ancient *D. punctatus* introgression in the Atlantic but decreased *D. punctatus* ancestry in the Mediterranean (Figure 2), thus strengthening the association of RI-islands to differential rates of archaic ancestry.

### Estimation of the time since introgression between D. punctatus *and* D. labrax

We estimated the timing of past gene flow between *D. punctatus* and *D. labrax* by first comparing the length distribution of *D. punctatus* tracts introgressed into Atlantic *D. labrax* genomes to that of Atlantic *D. labrax* tracts introgressed into western Mediterranean *D. labrax* genomes (Figure 3A). The two distributions showed similar shapes although *D. punctatus* tracts were on average almost ten-time shorter 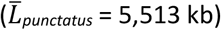 than Atlantic *D. labrax* tracts 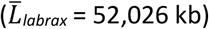. *D. punctatus* tracts were also less abundant in almost all length classes except for the shortest tracts (Figure 3A). We estimated 1 the average time since introgression for both distributions as 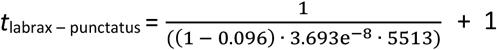 and 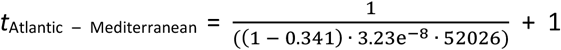, which placed the contact between *D. punctatus* and *D. labrax* approximately 6 times earlier than the one between the two *D. labrax* lineages. Using the age of secondary contact previously estimated between Atlantic and Mediterranean sea bass lineages (i.e. 11,500 years, Tine et al. 2014; Duranton et al. 2018) as a calibration time-point, ancient gene flow between the two species was dated to *ca.* 70,000 years ago. Secondly, we converted the estimated values of the transition parameter (*p*) of the HMM model used to detect archaic introgressed tracts to estimate one value of *T*_admix_ for each chromosome (Supplementary Table 2). From the obtained time distribution (Figure 3B), we estimated the most probable time of ancient admixture to *T*_admix_ = 91,149 (CI_90%_ = [85,831; 110,645]) years.

**Figure 3.**
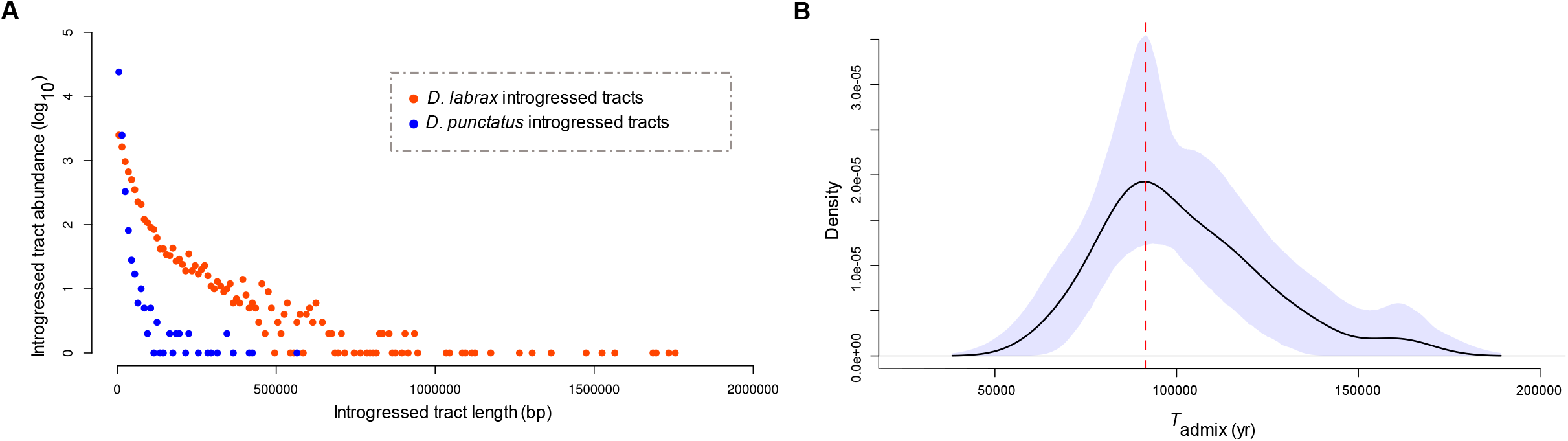
Estimation of the time since admixture between *D. punctatus* and Atlantic *D. labrax*. **A.** Length distributions of *D. punctatus* tracts introgressed into Atlantic *D. labrax* genomes (blue) and Atlantic *D. labrax* tracts introgressed into western Mediterranean *D. labrax* genomes (orange). Both distributions were generated using similar sequence lengths (totalizing 65.6 Mb) along the genomes of 14 Atlantic and 14 Mediterranean individuals, so that tract abundances can be compared. **B.** Distribution of estimated time since admixture between *D. punctatus* and *D. labrax* (*T*_admix_) obtained from estimated transition parameter values of the HMM model over the 24 chromosomes. The maximum of the distribution is represented by the vertical red dashed line and the blue shape represents the 95% credibility envelope of the distribution obtained using 10,000 bootstrap resampling.

### The frequency of *D. punctatus* derived mutations in *D. labrax*

We used the conditioned site frequency spectrum between Atlantic and Mediterranean lineages as a way to represent how derived *D. punctatus* alleles segregate in *D. labrax*. For SNPs that were not associated to RI-islands by the HMM approach, the one-dimensional distribution of allele frequencies (*CSFS*) was highly similar between Atlantic and Mediterranean *D. labrax* lineages, showing a bimodal shape with few intermediate frequency variants (Figure 4A). Most *D. punctatus* derived alleles were present at either low or high frequencies, with ancestral mutations almost fixed in both *D. labrax* lineages being about 100 times more abundant than *D. punctatus* derived mutations almost fixed in both *D. labrax* lineages in the *CJSFS* (Figure 4C). This result showed that the combined effects of incomplete lineage sorting and introgression during species divergence has resulted in very similar amounts of *D. punctatus* derived mutations between Atlantic and Mediterranean *D. labrax* lineages. By contrast, SNPs found to be associated with RI islands showed a large excess of *D. punctatus* derived alleles that were fixed or almost fixed in the Atlantic population, while segregating at low frequencies in the Mediterranean populations (Figure 4B and D). This remained true whatever the Mediterranean population (east, west or both) considered in the analysis (Supplementary Figure 7). The excess of high-frequency *D. punctatus* derived mutations in the Atlantic sea bass lineage was also clearly visible in the reversal of the *CSFS* in RI islands compared to non-RI regions (Figure 4A and B). Therefore, differential introgression of *D. punctatus* derived mutations in RI islands is most likely due to their direct role in reproductive isolation, rather than a delayed post-glacial rehomogenization due to already-existing genetic barriers between *D. labrax* lineages in these regions.

**Figure 4.**
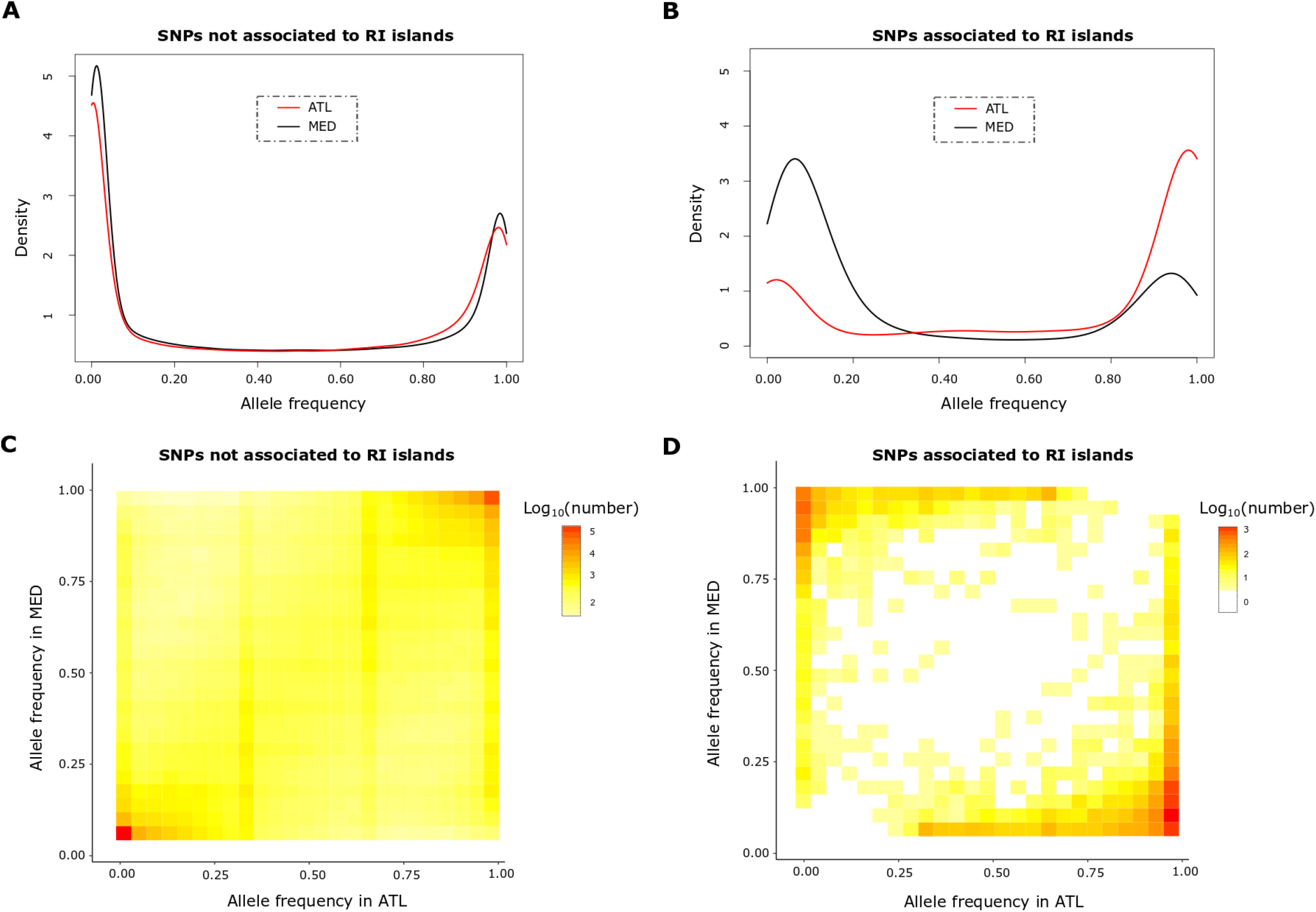
One and two-dimensional Site Frequency Spectra of *D. punctatus* derived alleles segregating in *D. labrax*. **A.** Conditional Site Frequency Spectra (*CSFS*) of *D. punctatus* derived alleles in AT (red) and MED (black) *D. labrax* lineages for categories of SNPs that are either not associated, or (**B**) associated to RI islands identified between the two *D. labrax* lineages. **C.** Conditional Joint Site Frequency Spectra (*CJSFS*) of derived *D. punctatus* alleles between MED (54 individuals) and AT (14 individuals) lineages based of 618,842 SNPs not involved in RI, and (**D**) 7,372 SNPs involved in RI.

## Discussion

Recent speciation genomics studies have revealed that genomic regions involved in RI often contain anciently diverged alleles <u>(e.g. Meier *et al.* 2017; Han *et al.* 2017; Nelson and Cresko 2018)</u>. One of the competing hypotheses to explain their origin is ancient admixture with an already diverged lineage. Our main objective here was to determine if such a scenario could explain the excess of divergence observed in RI regions between Atlantic and Mediterranean *D. labrax* lineages (30). To achieve this goal, we used different complementary approaches that collectively provided strong support for ancient introgression from the sister species *Dicentrarchus punctatus*. Despite low divergence (*d*_XY_ = 0.55%), partially overlapping range distributions and interfertility in artificial crosses (48), contemporary hybridization has not been observed in the wild between *D. labrax* and *D. punctatus* (Tine et al. 2014). We here show that interspecies admixture has likely happened earlier in the past, bringing new key elements to understand the complex evolutionary history of unachieved speciation between Atlantic and Mediterranean sea bass lineages.

### Extent of ancient admixture

Overall, the average fraction of contemporary genomes derived from ancient admixture was lower than 6% (i.e. 5.39% in the Atlantic and 2.82% in the Mediterranean lineage), which is only slightly higher than the estimated persistence of archaic ancestry in humans and brown bears (22,49). Whether these low background levels reflect a relatively limited contribution of genetic material from *D. punctatus* during admixture, or the impact of long-term selection against admixed foreign ancestry (29,50,51) was out of the scope of this study. Instead, we focused on understanding the marked excess of shared derived mutations found between *D. punctatus* and the Atlantic compared to the Mediterranean *D. labrax* lineage in RI-associated regions. This finding was strengthened by the locally increased frequency of archaic introgressed tracts found in Atlantic genomes within regions associated to RI with the Mediterranean lineage. Such locally elevated differences in the frequency of *D. punctatus* derived alleles explain the increased sequence divergence previously observed in RI islands between Atlantic and Mediterranean lineages (Duranton et al. 2018). Below, we consider potential limitations to the detection of archaic introgression from contemporary genomes, and the related challenge of dating ancient admixture. We then discuss how the genomic mosaicism of species ancestry may relate to different mechanisms potentially involved in European sea bass speciation.

### Separating ancient introgression from shared ancestral variation

Distinguishing past introgression and shared ancestral variation from ABBA-BABA and *f*_D_ statistics can be difficult, especially in regions of reduced divergence (39). Therefore, the positive correlations observed between *f*_D_, the inferred frequency of archaic segments, and *d*_XY_ provided good support that regions of high *D. punctatus* ancestry in the Atlantic are responsible for increased divergence between *D. labrax* lineages. Admittedly, past gene flow may also have occurred with another now extinct species rather than with *D. punctatus*, as it has been shown for other species (20,22,52). However, since *D. labrax* harbors shared derived alleles with *D. punctatus*, any alternative ghost donor lineage must have shared a long common history with the spotted sea bass.

Another potential issue with the tests performed to detect ancient admixture is that they often rely on differential introgression patterns between two candidate recipient populations (39,40). Therefore, these tests only enabled us to detect regions where the level of archaic introgression differs between Atlantic and Mediterranean *D. labrax* lineages. This problem could be particularly acute outside RI regions, where post-glacial gene flow between *D. labrax* lineages has almost completely rehomogenized allele frequencies (Tine et al. 2014; Duranton et al. 2018). To determine whether *D. punctatus* ancestry was simply absent or present but at similar levels in both lineages, we used the RND_min_ statistics that does not rely on the comparison of two populations (Rosenzweig et al. 2016). Low and nearly constant RND_min_ values indicated a widespread presence (although most of the time at low frequencies) of anciently introgressed tracts along Atlantic *D. labrax* genomes. By contrast, regions of elevated RND_min_ that coincided with the location of RI-islands revealed local resistance to introgression in Mediterranean *D. labrax* genomes. Therefore, both lineages contain *D. punctatus* introgressed tracts at relatively similar levels outside RI islands, which contrasys with strong archaic ancestry differences found within RI-islands.

### Timing of ancient introgression

To understand why *D. punctatus* alleles were rare within RI genomic regions in the Mediterranean, we reconstructed the history of ancient admixture by estimating the time of contact between *D. punctatus* and *D. labrax*. The two different methods respectively inferred a contact taking place approximately 70,000 and 90,000 years ago. Although these two estimates slightly differ, they both place ancient admixture during the last glacial period (53), when Atlantic and Mediterranean *D. labrax* lineages were inferred to be geographically isolated (30,32). The current distribution range of *D. punctatus* partially overlaps with the southern part of the *D. labrax* distributional area in both the Atlantic (i.e. from southern Biscay to Morocco) and southern Mediterranean Sea (i.e. North African shores). It is thus likely that the latitudinal range shifts that occurred during quaternary ice ages (54) have favored hybridization by further increasing the range overlap between the two species, as they were coexisting in the Iberian or the north-western African Atlantic refugium (55). Once the two *D. labrax* lineages came into secondary contact after the last glacial maximum, the *D. punctatus* alleles already introgressed within Atlantic genomes could have readily introgressed Mediterranean genomes. This hypothesis was supported by the observed gradient of decreasing *D. punctatus* ancestry from the Atlantic to the eastern Mediterranean lineage, which mirrored the gradient in Atlantic ancestry generated by the post-glacial secondary contact (30). The fact that *D. punctatus* tracts have most probably introgressed the Mediterranean lineage secondarily, indicates that ancient hybridization has only occurred in the Atlantic during the last glacial period. A possible explanation is the absence of sympatry between *D. punctatus* and *D. labrax* within the Mediterranean during the last glacial period. A missing piece of the reconstructed historical scenario remains with respect to the role of *D. punctatus* alleles in RI.

### Causative role of high-frequency *D. punctatus* alleles in RI-islands

If most of the currently observed RI-islands between *D. labrax* lineages were already existing before ancient admixture with *D. punctatus*, such genetic barriers would have impeded the introgression of *D. punctatus* alleles within the Mediterranean lineage (30). However, they would not account for increased frequencies of *D. punctatus* derived alleles within RI-islands in the Atlantic lineage. The fact that, in Atlantic *D. labrax*, regions associated to RI exhibited closely fixed *D. punctatus* derived alleles that comparatively occurred at low frequencies elsewhere in the genome strongly supports their direct role in the establishment of RI. This finding thus indicates that *D. punctatus* alleles have been first locally driven to high frequencies in the Atlantic *D. labrax* lineage, while being secondarily prevented from introgression within the Mediterranean lineage.

### Why anciently introgressed alleles contribute to RI?

#### Locally adaptive introgression

Understanding the underlying evolutionary mechanisms through which admixture has contributed to the buildup of reproductive isolation remains highly challenging (56,57). One evolutionary force that can drive an allele to fixation is local positive selection. *D. punctatus* alleles may have fixed in the Atlantic *D. labrax* lineage following admixture because they provided a selective advantage in the Atlantic environment compared to ancestral *D. labrax* alleles, a process called adaptive introgression (58). Several studies have revealed that the acquisition of adaptive phenotypes can be done through hybridization, such as altitude adaptation in humans (59), mimicry in *Heliconius* butterflies (60) or among others, seasonal camouflage in the snowshoe hares (61). Indeed, adaptive introgression allows the rapid transfer of linked variants that have already been tested by natural selection in their original environment, thus facilitating local adaptation (62). Therefore, it is theoretically possible that the Atlantic *D. labrax* lineage has received from *D. punctatus* advantageous alleles in the Atlantic environment that revealed to be deleterious in the Mediterranean Sea. Nevertheless, adaptive introgression is usually difficult to prove since it can be confounded with other processes such as uncoupling of an incompatibility from a multilocus genetic barrier (Fraïsse *et al.* 2014). Furthermore, it has been argued that adaptive introgression cannot play an important role in reproductive isolation, because unconditionally favorable alleles spread easily between diverging lineages until RI is nearly complete (65).

#### Fixation-compensation of deleterious mutations

Another evolutionary force that may have driven *D. punctatus* derived alleles to fixation is genetic drift, which can induce the fixation of deleterious mutations and thus increase mutation load (66). When gene flow occurred between *D. punctatus* and *D. labrax* during the last glacial period, populations of each species were probably experiencing bottlenecks (54), which decreased the efficiency of selection and enhanced the probability to fix deleterious mutations by drift. Weakly deleterious *D. punctatus* alleles may therefore have introgressed and fixed within the *D. labrax* Atlantic population. Another related mechanism that may have influenced the outcome of hybridization is associative overdominance, due to the masking of recessive deleterious mutations in admixed genotypes (Whitlock et al. 2000; Bierne *et al.* 2002). Heterosis can locally increase the introgression rate of foreign alleles, even if interbreeding populations have similar amounts of deleterious variation (68). Therefore, heterosis may have favored the introgression of weakly deleterious *D. punctatus* variants in a bottlenecked Atlantic *D. labrax* lineage. Subsequently, when Atlantic and Mediterranean *D. labrax* lineages reconnected following postglacial recolonizations, expanding populations would have been sufficiently large to reveal the deleterious effects of the introgressed alleles, generating hybrid depression and hybridization load (29,69). Furthermore, the Atlantic population may have had enough time to evolve compensatory mutations (70), which could have become substrate for increased RI. The fact that most genomic regions involved in RI between *D. labrax* lineages exhibit low recombination rates (Tine *et al.* 2014; Duranton et al. 2018) could indicate a role of slightly deleterious alleles in RI, since selection is less efficient when linkage is strong.

#### Reciprocal sorting of DMIs

Reproductive isolation may also have evolved through the resolution of genetic conflicts resulting from the contact between two diverged populations (71,72). Because each population has almost inevitably fixed new adaptive or nearly neutral variants that reveal incompatible when combined in hybrid genomes (73), Bateson-Dobzhansky-Muller incompatibilities (BDMIs) are recognized as a common substrate for speciation (2). A genomic conflict induced by a two-locus BDMI can be resolved by fixing one of either parental alleles. In a hybrid population generated by an equal mixture of individuals from both parental populations, there is a 50% chance of fixing either parental combination (71). Therefore, the resolution of multiple BDMIs in an admixed population offers ample opportunity to reciprocally resolve independent BDMIs with respect to the origin of the parental allelic combination, which results in RI from both parental populations. Even in the presence of skewed initial admixture proportions, fixation of the minor parent combination can still happen with a sufficient number of BDMIs (71). Therefore, the resolution of genetic conflicts between *D. punctatus* and *D. labrax* alleles in the Atlantic lineage may have induced the fixation of *D. punctatus* alleles at some incompatibility loci. Upon contact between Atlantic and Mediterranean *D. labrax* lineages, fixed *D. punctatus* alleles may have recreated the BDMIs, thus contributing to RI. This non-adaptive speciation model due to selection against genetic incompatibilities has the advantage to explain both the fixation of *D. punctatus* alleles within the *D. labrax* Atlantic population, and their incompatibility with the Mediterranean lineage. Verbally, it can be seen as a case whereby speciation reversal between lineages A and B contributes to strengthen RI between lineages B and C through the transfer of incompatibilities between two porous species boundaries.

### Conclusion

To conclude, our results show that divergent haplotypes that were introgressed from *D. punctatus* about 80,000 year ago have contributed to the strengthening of nascent RI between Atlantic and Mediterranean *D. labrax* lineages. The resulting genomic architecture of RI between contemporary *D. labrax* lineages is thus constituted by a mosaic of fixed blocks of different ancestries, that is, a mixture of genetic barriers inherited from the own *D. labrax* divergence history and the contribution of ancient admixture. Although additional analyses will be needed to fully understand which process has driven the fixation of *D. punctatus* alleles within Atlantic genomes, the resolution of genetic conflicts between *D. punctatus* and *D. labrax* seems the most parsimonious hypothesis (Schumer *et al.* 2015; Blanckaert and Bank 2018). This speciation mechanism can be thought of as a transfer of incompatibilities between two species boundaries, from the strongest to the weakest barrier, which is eventually strengthened by the displacement of genetic conflicts inherited from an ancient episode of admixture. Our finding adds to previous reports showing that postglacial and recent hybridization events have played a role in the buildup of RI between admixed and parental lineages by generating similar genomic mosaics of ancestries (29,74,75). The contribution of ancient admixture in European sea bass speciation suggests that significantly older admixture events, which may have left cryptic signatures in contemporary genomes, can be involved in seemingly recent speciation histories.

## Material and methods

### Whole-genome resequencing and haplotyping

We sequenced the whole genome of one *Dicentrarchus punctatus* individual from the Atlantic Ocean (Gulf of Cadiz, PUN) and 59 new *Dicentrarchus labrax* individual genomes. Fifty-two of them were wild individuals captured from the Atlantic Ocean (English Channel, 10 males ♂_AT_), the western Mediterranean Sea (Gulf of Lion, 14 females ♀_WME_ and 9 males ♂_WME_) and the eastern Mediterranean Sea (Turkey, 10 males ♂_NEM_ and Egypt, 9 males ♂_SEM_). Some of these specimens were involved in experimental crosses to generate first generation hybrids. Seven F1 hybrids obtained from 7 different biparental families (pedigree ♂_AT_ x ♀_WME_) were also submitted to whole-genome sequencing. All captive breeding procedures were performed at Ifremer’s experimental aquaculture facility (agreement for experiments with animals: C 34-192-6), where fish were reared in normal aquaculture conditions in agreement with the French decree no. 2013-118 (1 February 2013 NOR:AGRG1231951D).

Whole genome sequencing libraries were prepared separately for each individual using either the Illumina TruSeq DNA PCR-Free (40 individuals) or the TruSeq DNA Nano protocol (20 individuals), depending on DNA concentration (Supplementary Table 1). Pools of 5 individually barcoded libraries were then sequenced on 12 separate lanes of an Illumina HiSeq3000 using 2×150bp PE reads at the GeT-PlaGe Genomics platform (Toulouse, France). Thirty-three individuals were sequenced twice due to insufficient amounts of sequence reads obtained in the first run (Supplementary Table 1). For each individual, the alignment of PE reads to the sea bass reference genome (32) was performed using BWA-mem v0.7.5a (33) with default parameters. Duplicate reads were marked using Picard version 1.112 before being removed, producing a mean coverage depth of 33.8X per individual (Supplementary Figure 1). We then followed GATK’s (version 3.3-0-g37228af) best practice pipeline for individual variant calling (using HaplotypeCaller), to joint genotyping, genotype refinement and variant filtering (using Filter Expression: QD<10; MQ<50; FS>7; MQRankSum<-1.5; ReadPosRankSum<-1.5). We used the BQSR algorithm to recalibrate base quality scores using a set of high-quality variants identified in a previous study (30), and to perform variant quality score recalibration using the VQSR algorithm. Hard filtering was then applied to exclude low-quality genotypes with a GQ score < 30. For the 7 mother-father-offspring trios, we used family-based priors for genotype refinement. We obtained a total of 14,579,961 SNPs after filtering for indels, missing data (using --max-missing-count 8) and removing the mitochondrial and ungrouped scaffolds (chromosome UN) in VCFtools v0.1.11 (34).

We performed haplotype phasing in *D. labrax* after removing the *D. punctatus* individual and merging the 59 newly sequenced genomes with the 16 genomes already obtained in Duranton *et al.* (2018). Fifteen individuals that were involved in family crosses (i.e. newly sequenced or not already phased in the previous study) were submitted to phasing-by-transmission using the PhaseByTransmission algorithm in GATK with default parameters and a mutation rate prior of 10^−8^ for *de novo* mutations. For all individuals, variants located on a same read pair were directly phased using physical phasing information. Non-related *D. labrax* individuals were then statistically phased using the reference-based phasing algorithm implemented in Eagle2 (version 2.4) (35). The 22 parents phased with the phasing-by-transmission approach were used to build a European sea bass reference haplotype library (F1 genomes were excluded since their haplotype information was redundant with that of their parents), which was used in Eagle2 to improve statistical phasing. We finally filtered out SNPs that were not phased or not genotyped over all individuals (using --max-missing-count 0 and – phased in VCFtools), to generate a dataset of haplotype-resolved whole-genome sequences from 68 unrelated *D. labrax* individuals (14 AT, 31 WME, 11 SEM and 12 NEM), containing 5,074,249 phased SNPs without missing data. The genetic relationships of the newly sequenced genomes with respect to the 16 already available was evaluated with a Principal Component Analysis (Supplementary Figure 2). Although we detected a slight genetic differentiation between North and South eastern Mediterranean samples on the PCA (Supplementary Figure 2), we later determined that they present similar genome-wide average levels of Atlantic ancestry and introgressed tract length (18,148 bp for the South and 17,769 bp for the North, Supplementary Figure 5). Therefore, we regrouped these samples together within a single eastern Mediterranean population, similarly to Duranton et al. (2018).

### Phylogenomic analyses

We used RAxML v.8.2.12 (36) to generate maximum-likelihood trees of Moronids genomes in non-overlapping 50 kb windows (including *Morone saxatilis*, *D. punctatus* and the Atlantic and Mediterranean *D. labrax* lineages). Ancient admixture is expected to generate discordant trees among genomic windows. However, if admixture is ancient, introgressed tracts may be too short to influence the phylogenetic signal in 50 kb windows. Therefore, we did the same analyses using a 2 kb window size to increase the resolution of local genealogies while keeping enough informative sites. In order to account for disparities among species’ genome sequence datasets, we used only one individual haplome for each species/lineage for this analysis. The alignment of these four haplomes spanned 52% of the *D. labrax* genome, a fact largely due to the fragmentation of the *M. saxatilis* (SAX) genome that produced discontinuous local alignments to the *D. labrax* reference genome (30). In order to account for this fragmentation, we only analyzed windows with less than 10% missing data in local alignments. We obtained 3,329 and 155,155 trees under the GTRGAMMA model for analyses based on 50 and 2 kb windows, respectively. Trees generated in windows of similar size were then superposed using DensiTree v2.2.5 (37) for visualization. In order to provide indications for genome-wide average absolute sequence divergence between all pairs of species and lineages, we calculated *d*_XY_with the same individual haplomes used for the RAxML analysis using MVFTools v5.1.2 (38) and averaged distance values calculated in non-overlapping 50kb windows.

### Tests for foreign introgression within *D. labrax*

We tested for admixture between *D. labrax* and another species using three different methods that capture complementary aspects of the data. Since *D. punctatus* is the only closely related species parapatrically distributed with *D. labrax*, we first tested for historical gene flow between these two species. To do so, we used the ABBA-BABA test (19,23) with *M. saxatilis* as the outgroup (O), *D. punctatus* as the potential donor species (P3) and the two *D. labrax* lineages as potential recipient populations (P1 and P2). We used the dataset containing 14,579,961 SNPs from *D. punctatus* and unphased *D. labrax* samples, and only kept sites that were available for both *M. saxatilis* and *D. punctatus* in genome alignments, representing a total of 9,606,462 SNPs. This allowed testing for different amounts of gene flow between P3 and P2, and P3 and P1, by comparing the number of genealogies of type ((P1,(P2,P3),O) (*i.e.* ABBA genealogies) and ((P2,(P1,P3),O) (*i.e.* BABA genealogies). An excess of shared derived alleles between the donor and one of the two recipient populations (i.e. excess of ABBA over BABA genealogies, or vice versa) indicates gene flow from *D. punctatus* to *D. labrax* population P2 or P1, respectively. Although the ABBA-BABA test is adequate to detect introgression, the Patterson’s *D* statistic that measures the imbalance between the two types of genealogies 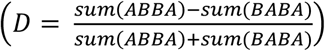 is not appropriate to quantify introgression over small genomic windows (39). Therefore, we used the *f*_D_ statistics 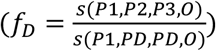 to estimate admixture proportion between P2 et P3, were *s* = sum(ABBA-BABA) and *PD* corresponds to the most likely donor population (i.e. the population with the higher frequency of the derived allele). In order to test for admixture between *D. punctatus* and different populations of *D. labrax*, we made different tests using successively the Atlantic (AT), eastern (EME combining SEM and NEM individuals), western (WME) or the whole Mediterranean (MED) populations of *D. labrax* populations as P2 or P1. We used scripts from Martin *et al.* (2015) to estimate the number of ABBA and BABA genealogies and the *f*_D_ statistics in non-overlapping 50 kb windows along the genome, keeping only windows containing at least 500 SNPs.

Secondly, we used a method that allows identifying archaic introgressed tracts without using an archaic reference genome for the donor species (40). The main advantage of this method is that it makes no assumption on the identity of the donor species. Basically, it looks for local excesses of private variants in a candidate recipient population by comparison to another non-admixed population (40). In order to test for archaic introgression within the Atlantic *D. labrax* lineage, we identified variants that were not shared with the eastern Mediterranean population, and conversely to test for introgression in the Mediterranean *D. labrax* lineage. We only analyzed the eastern Mediterranean population because the western Mediterranean is more strongly impacted by gene flow from the Atlantic (30,32). We used the phased genomes dataset containing 5,074,249 SNPs, assuming a constant mutation and call rate to run the model in 1000 bp windows along each chromosome. For each window, the probability that an individual haplotype contains an archaic introgressed fragment was estimated to identify introgressed windows with a posterior probability superior to 0.8 (40). We then combined individual profiles of introgressed windows to estimate the fraction of introgressed archaic tracts in each population (*F*_archaic_), as the fraction of haplotypes for which a window was identified as introgressed. The inferred fraction of introgressed archaic tracts was finally averaged in non-overlapping 50 kb windows along the genome.

Finally, we used the RND_min_ statistics, which is sensitive to rare introgression while being robust to mutation rate variation across the genome (41). The main advantage of this statistic is that, unlike the two former methods, it does not rely on the comparison of two recipient populations that differ in their level of introgression. The RND_min_ corresponds to the ratio of the minimal pairwise distance between haplotypes from the potential donor and recipient populations (*d*_min_) over the average divergence of those populations to an outgroup species (*d*_out_). If gene flow has occurred genome-wide, then locally elevated RND_min_ values indicate regions where introgression has been limited or absent. We used MVFTools v5.1.2 (38) to measure *d*_min_ between *D. punctatus* and different population of *D. labrax* (AT, EME, WME, MED). For this analysis, we used 4,943,488 SNPs polymorphic sites that were phased within *D. labrax* and non-missing in *D. punctatus*. On one hand, this dataset excludes a large number of variants that are differentially fixed between *D. punctatus* and *D. labrax*, and therefore underestimates the real level of divergence between *D. punctatus* and *D. labrax.* On the other hand, excluding diagnostic SNPs rendered the test more sensitive to the detection of ancient introgression, since the accumulation of divergence after introgression only adds noise to chromosomal variations in RND_min_. We estimated *d*_out_ by averaging the divergence measures between *M. saxatilis* and the two *Dicentrarchus* species. All values were averaged in non-overlapping 50 kb windows along the genome.

### Detection of introgressed tracts between Atlantic and Mediterranean *D. labrax* lineages

In order to test whether ancient introgression has influenced genomic patterns of post-glacial gene flow between Atlantic and Mediterranean *D. labrax* lineages, we mapped Atlantic tracts introgressed into Mediterranean genomes and conversely. Local ancestry inference was performed with Chromopainter v0.04 (42), an HMM-based program that estimates the probability of Atlantic and Mediterranean ancestry for each variable position along each haplome. To do so, it compares a focal haplotype to reference populations composed of non-introgressed Atlantic and Mediterranean haplotypes. Since Mediterranean individuals are introgressed to various extents by Atlantic alleles, we used a pure Mediterranean reference population reconstituted by Duranton *et al.* (2018) with the same model parameters. We then identified the starting and ending position of each introgressed tract within both Atlantic and Mediterranean genetic backgrounds by analyzing the ancestry probability profiles inferred by Chromopainter, following the same methodology as in Duranton *et al.* (2018). Identified tracts in each *D. labrax* population (Supplementary Figure 5) were then combined to estimate the fraction of introgressed tracts (*F*_intro_) for each position along the genome, which was finally averaged in non-overlapping 50 kb windows.

### Delineation of RI regions between Atlantic and Mediterranean *D. labrax* lineages

We adapted the HMM approach developed by Hofer *et al.* (2012) to precisely delineate genomic regions involved in RI between Atlantic and Mediterranean *D. labrax* lineages. Genomic regions involved in RI between European sea bass lineages are characterized by elevated genetic differentiation and increased resistance to gene flow (30,32). Therefore, we combined both measures of *F*_ST_ (43,44) and resistance to introgression measured as the inverse of *F*_intro_ (28,30). To identify true RI islands in our HMM strategy, we thus used the ratio of *F*_ST_ over *F*_intro_ (i.e. the frequency of Atlantic tracts within western Mediterranean *D. labrax* genomes). Our rationale was that these regions should be associated with both high *F*_ST_ (Supplementary Figure 6A and D) and low *F*_intro_ values (Supplementary Figure 6B and E) (30), hence elevated *F*_ST_/*F*_intro_ ratio values (Supplementary Figure 6C and F). We used the HMM approach to map RI at two different scales, a SNP-by-SNP (Supplementary Figure 6A-C) and a 50kb window scale, which was more suitable to delineate regions (Supplementary Figure 6D-F). We used VCFtools v0.1.15 (34) to estimate *F*_ST_ between the Atlantic and the western Mediterranean *D. labrax* lineage for each SNPs and every non-overlapping 50kb window along the genome. The HMM was designed with three different states corresponding to low (i.e. neutral genomic regions), intermediate (i.e. regions experiencing linked selection) and high *F*_ST_/*F*_intro_ ratio values (i.e. regions involved in RI). The most likely state of each SNP/window was inferred by running the HMM algorithm chromosome by chromosome. Finally, we controlled for false discovery rate and retained only SNPs/windows with an FDR-corrected p-value inferior to 0.001 (43).

### Estimation of the time since foreign introgression within *D. labrax*

We used two different approaches to estimate the time since foreign introgression within *D. labrax*. First, we relied on the fact that the length of introgressed tracts is informative of the time elapsed since introgression. Recombination progressively shortens migrant tracts across generations following introgression into a new genetic background (45,46). Since we found a good correspondence between the inferred fraction of introgressed archaic tracts (*F*_intro-archaic_) and *f*_D_ values (see results) using *D. punctatus* as a donor species, we used the length of archaic haplotypes that were identified with the method of Skov *et al.* (2018). Only archaic tracts found in Atlantic genomes within windows involved in RI between Atlantic and Mediterranean *D. labrax* lineages were considered, corresponding to 1310 windows of 50 kb. This choice was justified because archaic tract detection relies on a signal of differential introgression between two populations. Therefore, archaic tracts can be correctly identified and delimited only if they are present in one lineage (e.g. the Atlantic) but absent in the other (e.g. the Mediterranean), which was only the case in RI islands between Atlantic and Mediterranean *D. labrax* lineages (see results).

Under simple assumptions, there is an analytical expectation for the average length of introgressed tracts 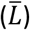 as a function of the number of generations since introgression (*t*), the local recombination rate (*r* in Morgans per bp) and the proportion of admixture (*f*), which takes the form 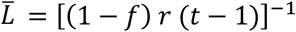 (26). We used this equation to estimate the age of admixture between *D. punctatus* and the Atlantic lineage of *D. labrax* (*t*_labrax – punctatus_), as well as between the two lineages of *D. labrax* (*t*_Atlantic – Mediterranean_). For each estimation, we used the average value of the retained windows. Hence, for *t*_labrax – punctatus_: *f* = 0.096, *r* = 3.693e^−8^ M/bp (32) and 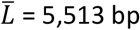, and for *t*_Atlantic – Mediterranean_: *f* = 0.341, *r* = 3.23e^−8^ M/bp, and 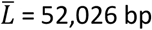. Since we only considered a relatively small fraction of the genome to call archaic tracts, we could not obtain precise estimations of those parameters. Therefore, we estimated the age of contact between *D. punctatus* and *D. labrax* by reference to the age of the post-glacial secondary contact between Atlantic and Mediterranean lineages of *D. labrax*, which has been more precisely estimated to 2,300 generations using a larger fraction of the genome (30,32).

Secondly, we transformed the estimated transition parameter values of the HMM model used to detect archaic introgressed tracts using *p* ≈ *T*_*admix*_ ⋅ 2 ⋅ *r* ⋅ *L* ⋅ *a* (40). In this equation, *p* is the probability of transition from the *D. labrax* to the archaic ancestry state, *T*_*admix*_ represents the admixture time in generations, *r* the recombination rate in Morgan per bp, *a* the admixture proportion and *L* the size of the window (here *L* = 1000 bp). Parameter *p* was estimated separately for each chromosome by averaging over the values estimated per individual haplome. We finally estimated *T*_*admix*_ chromosome by chromosome using the average recombination rate and the fraction of archaic introgressed tracts of each chromosome (Supplementary Table 2). The time in generations was converted into years using a generation time of 5 years (32). We then obtained a distribution for *T*_*admix*_ across the 24 chromosomes, from which we identified the maximum and its 90% confidence interval by bootstrapping the distribution 10,000 times.

### Characterizing foreign ancestry tracts within *D. labrax*

We used Spearman’s correlation test to evaluate relationships among *F*_intro-archaic_, *f*_D_, *d*_XY_ and RND_min_ statistics that relate to a series of predictive hypotheses. More specifically, if *D. punctatus* has anciently contributed to *D. labrax* in the Atlantic, the local abundance of archaic tracts inferred within Atlantic *D. labrax* genomes (*F*_archaic_) should be positively correlated to *f*_D_. Moreover, if the abundance of archaic tracts within Atlantic *D. labrax* explains the presence of anciently diverged alleles between Atlantic and Mediterranean sea bass lineages, *d*_XY_ should increase with the amount of ancient admixture. Finally, if regions involved in RI between Atlantic and Mediterranean sea bass lineages harbor reduced frequencies of *D. punctatus* derived tracts in the Mediterranean, a positive correlation is expected between RND_min_ measured between *D. punctatus* and Mediterranean *D. labrax* and ancient admixture from *D. punctatus* within the Atlantic.

We then focused on SNP-level statistics to specifically address the frequency distributions of derived mutations from *D. punctatus* within *D. labrax* genomes, separately in the Atlantic and Mediterranean lineages. Since anciently introgressed alleles most likely originated from *D. punctatus* (see results), we used *D. labrax* polymorphic sites for which *M. saxatilis* harbors the ancestral and *D. punctatus* the derived state (*i.e.* ABBA-BABA informative sites) to characterize ancient introgression. For each of these SNPs, we measured the frequency of the *D. punctatus* derived allele separately in the Atlantic and Mediterranean *D. labrax* populations using VCFtools. We then separated SNPs associated to RI islands from those that were not associated to RI islands in the SNP-based HMM analysis to represent the site frequency spectrum of each *D. labrax* lineage, conditioned on *D. punctatus* being derived (*CSFS*). Finally, two conditioned joint site frequency spectra (*CJSFS*) were generated (i.e. for RI and non-RI SNPs) to represent the bi-dimensional SFS between Atlantic and Mediterranean *D. labrax* lineages, conditioning on sites that have the derived allele in *D. punctatus*. These analyses aimed at distinguishing two hypotheses with respect to the mechanisms underlying differential introgression of *D. punctatus* derived mutations in RI islands between Atlantic and Mediterranean *D. labrax*. Our first hypothesis was that anciently introgressed alleles are not directly involved in RI but simply maintained at different frequencies because genetic barriers between *D. labrax* lineages (i.e. unrelated to the history of ancient admixture) have impeded their post-glacial rehomogenization. In this case, we expected no excess of high-frequency *D. punctatus* derived mutations in RI islands compared to non-RI regions. Alternatively, under the hypothesis that anciently introgressed alleles are associated with reproductive isolation in sea bass, an excess of *D. punctatus* derived mutations almost fixed within RI islands in the Atlantic but nearly absent in the Mediterranean was expected compared to the alternate configuration (i.e. almost fixed in the Mediterranean and nearly absent in the Atlantic).

## Ackowledgement

This work was co-founded by the GeneSea project (n° R FEA 4700 16 FA 100 0005) by the French Government and the European Union (EMFF, European Maritime and Fisheries Fund) at the “Appels à projets Innovants” managed by the FranceAgrimer Office and the ANR grant CoGeDiv (ANR-17-CE02-0006-01 to P.-A.G). This work was performed in collaboration with the GeT core facility, Toulouse, France (http://get.genotoul.fr), and was supported by France Génomique National infrastructure, funded as part of “Investissement d’avenir” program managed by Agence Nationale pour la Recherche (contract ANR-10-INBS-09). We also thank Ifremer’s experimental aquaculture platform for the breeding and the rearing of the hybrid populations

## Supporting Information Legends

**Supplementary Table 1** – Summary statistics of sequencing and mapping data for each individual. Individuals whose name is in bold are those involved in crossing.

**Supplementary Figure 1** - **Depth of coverage per individual**. Median (dark gray), first (light gray) and third (black) quartile of the depth of coverage for the 10 Atlantic males (AT), the 23 individuals from the western Mediterranean sea (14 females (F) and 9 males (WM/WME)), the 19 individuals from the eastern Mediterranean sea (9 males from the south (SEM) and 9 males from the north (NEM)), the 7 hybrids (F1) and the D. punctatus individual (Punc).

**Supplementary Figure 2** - Principal Component Analysis of the European sea bass population genetic structure. The analysis was performed on the 52 newly sequenced genomes (colored circles) and the 16 from a previous data set (1) (gray circles with colorful outline). We used the R package adegenet (2) on 91,073 SNPs with a minor allele frequency > 0.4. Individuals originated from four different geographic locations the Atlantic ocean (red, AT), the west (orange, WME), the north-east (dark yellow, NEM) and the south east (light yellow, SEM) of the Mediterranean sea. The first PCA axis explains 39.76% of the total inertia and distinguish the Atlantic and Mediterranean populations while the second PCA axis explains 6.55% of the total inertia and reveals a structure within the Mediterranean population.

**Supplementary Figure 3** –Consensus trees of the 155,155 Maximum-likelihood trees inferred in 2kb windows along the genome between M. saxatilis, D. punctatus and Atlantic and Mediterranean D. labrax lineages. There were four different topologies, the most frequent representing the species tree; 94.5% (blue), the second one grouping the Atlantic lineage with D. punctatus; 2.87 % (orange), a third one grouping D. punctatus with the Mediterranean lineage; 1.68 % (green) and a last one with unresolved relationship between D. labrax lineages and D. punctatus; 0.05% (not showed).

**Supplementary Figure 4** – Statistics measured in non-overlapping 50 kb windows along the genome. A. dXY measured between the Atlantic and Mediterranean (including eastern and western population) D. labrax lineage B. fD statistic measured using in red (((MED, AT), PUN), SAX), in green (((AT, WEST), PUN), SAX) and in blue (((AT, EAST), PUN), SAX). C. Fraction of archaic introgressed tracts (Farchaic) in the eastern Mediterranean and Atlantic population of D. labrax. D. RNDmin measured between D. punctatus and D. labrax Atlantic (red), western (green) and eastern (blue) Mediterranean populations. E. Ratio of FST and Fintro used to run the HMM approaches on 50 kb windows that rely on 3 states 1 (light grey), 2 (medium grey) and 3 (dark grey). Red points passed the control for false discovery. We defined island of reproductive isolation as continuous regions containing only red and dark grey points (red boxes). F. Ratio of FST and Fintro used to run the HMM approaches on SNPs, purple points are SNPs identified as involved in reproductive isolation that passed the control for false discovery.

**Supplementary Figure 5** – Introgressed tract length distributions. Length distributions of Mediterranean tracts introgressed into the Atlantic population (red) and of Atlantic tracts introgressed into the western (orange), north-eastern (dark yellow) and south-eastern (light yellow) Mediterranean populations. Distributions were generated over the whole genome using 11 individuals per population.

**Supplementary Figure 6** – Data and results for the SNPs and 50kb window based HMM approach to identify regions involved in reproductive isolation between the two lineage of D. labrax along chromosome 7. A. FST measured between the Atlantic and western Mediterranean population of D. labrax for each SNPs and in every non-overlapping 50 kb windows (D). B. Fraction of Atlantic tracts introgressed in western Mediterranean genomes (Fintro) for each SNPs and in every non-overlapping 50 kb windows (E). C. Statistic analyzed by the HMM approaches (FST divided by Fintro) for each SNPs and every 50 kb non-overlapping window (F). Ratio of FST and Fintro used to run the HMM approaches that rely on 3 states that identify; neutral genomic regions (state 1, light grey), genomic regions under linked selection (state 2, medium grey) and genomic regions involved in reproductive isolation (state3, dark grey). Red points are those that passed the control for false discovery. For the window approach we defined island of RI as continuous regions containing only red and dark grey points (red boxes).

**Supplementary Figure 7** – Distributions and joint allele-frequency spectrums of derived D. punctatus alleles present in D. labrax. Distribution of D. punctatus derived alleles frequency in AT (red) and WEST (black) (A) or East (D) D. labrax individuals for loci involved (solid line) or not (dashed lines) in reproductive isolation between the two D. labrax lineages. B. Joint allele-frequency spectrum of derived D. punctatus allele for the WEST (31 individuals) and AT (14 individuals) populations for 594,797 SNPs not involved and 7,372 SNPs involved (C) in reproductive isolation. E. Joint allele-frequency spectrum of derived D. punctatus allele for the EAST (23 individuals) and AT (14 individuals) populations for 594,454 SNPs not involved and 7,366 SNPs involved (C) in reproductive isolation.

**Supplementary Table 2** – values used to estimate Tadmix for each chromosome.

